# Transcriptomic Characterization of Soybean Roots in Response to *Bradyrhizobium* Infection by RNA Sequencing

**DOI:** 10.1101/024224

**Authors:** Qingyuan He, Yang Hongyan, Ping Wu, Zhengpeng Li, Songhua Wang, Changwei Zhu

## Abstract

Legumes interact with *rhizobium* convert N_2_ into ammonia for plant use. To investigate the plant basal nitrogen fixation mechanisms induced in response to *Bradyrhizobium*, differential gene expression in root of inoculated and mock-inoculated soybean was analysed by RNA-Seq. A total of 55787 transcripts were aligned to soybean genome reference sequences, 280 and 316 transcripts were found to be up-and down-regulated, respectively, in inoculated relative to mock-inoculated soybean’s root at V1 stage. Gene ontology (GO) analyses detected 104, 182 and 178 genes associated with cell component category, molecular function category and biological process category, respectively. Pathway analysis revealed that 98 differentiallly expressed genes (115 transcripts) involved in 169 biological pathways. We selected 19 differentially expressed genes and analyzed their expressions in mock-inoculated, inoculated USDA110 and CCBAU45436 using qRT-PCR. The results were consistent with those obtained from *rhizobia* infected RNA-Seq data. These showed that the results of RNA-Seq have reliability and universality. Additionally, this study showed some novel genes associated with nitrogen fixation process comparison with the previously identified QTLs.

---

Nitrogen is the most limiting element for crop growth and usually supplied by application of fertilizer, which brings on substantial costs to farmers and with potentially adverse effects on the environment. The leguminous plants establish a symbiotic relationship with *rhizobia* (symbiotic nitrogen fixation, SNF) to directly capture N_2_ to support plant growth. Nitrogen conversion takes place in a unique organ (root nodule). The development of root nodules commences with a molecular dialogue between the host plant and a compatible strain of *rhizobium*, involving a succession of complex process that lead to profound changes in both symbionts (Oldroyd *et al.*, 2011).

The plant excretes molecular signals, flavonoids, phenolic compounds which induce the synthesis of specific *rhizobia*-produced lipo-chito-oligosaccharides, called Nod factors (NFs). NFs directly stimulate their putative receptor protein (NFRs), such as LjNFR1/5 (Nod factor through receptor-like kinase) in *Lotus japonicus* and MtLYK3 (LysM-receptor like kinas 3)/ NFP (Nod factor perception) in *Medicago Trucatula* (Arrighi *et al.*, 2006; Limpens *et al.*, 2003; Madsen *et al.*, 2003; Radutoiu *et al.*, 2003), which are LysM (peptidoglycanbinding lysine motif) receptor-like of the legumes (Kochi *et al.*, 2010; Subramanian *et al.*, 2006). Many molecular events are triggered in a coordinate manner, leading to morphological and physiological changes in the host plant, necessary for a successful symbiosis (Oldroyd *et al.*, 2011). Bacterial attachment to the root hair and induces a calcium influx and membrane depolarization, subsequently lead to deformation and root fair curling, forming infection thread. Concomitantly, certain cortical cells divide to form nodule primorda, further development gives rise to nodules that differ from tumours in having defined anatomical structures. *Rhizobia* are ramified into these tissues, with subsequent release to the bacterium into plant cells where they differentiate into bacteriods and begin to fix nitrogen (Day *et al.*, 2000; Irving *et al.*, 2000).

Nodule formation and accommodation of endosymbiotic *rhizobia* inside nodules are strictly controlled by host plant genes. Plant genes that show enhanced expression during nodulation are named “nodulins” (Bladergron and Spaink, 1998), Such as ENOD2, ENOD12, ENOD40 (Kochi and Hata, 1995; Pichon *et al.*, 1992; Papadopoulou *et al.*, 1996). Several genes have been identified that play hinge functions in the perception and transduction of the bacterial NFs (Oldroyd and Downie, 2006, 2008; Jones *et al.*, 2007). NFRs are activated by NFs and subsequent stimulation of downstream signaling pathways through nuclear Ca^2+^ spiking. A perinuclear-anchored cation channel, MtDMI1 (Does-not-make-infections 1)/ LjCASTOR/ LjPOLLUX plays a critical role for upstream of Ca^2+^ spiking during early *rhizobia* infection, and its function appears to be regulated by an upstream component, MtDMI2/ LjSYMRK (Symbiosis receptor kinase)/MsNORK (Nodulation receptor kinase), which is a member of the LRR-RLK family (Leucine rich repeat-receptor like kinase) (Catoira *et al.*, 2000; Endre *et al.*, 2002; Madsen *et al.*, 2003). DMI1 and DMI2 act downstream of NFRs in the Nod factor signaling pathway (Radutoiu *et al.*, 2003). Ca^2+^-related MtDMI3 is a nuclear-localized CCaMK (Ca^2+^-calmodulin-dependent kinase) that functions downstream of Ca^2+^ spiking (Tirichine *et al.*, 2006). Activated DMI3 directly interacts with and activates nodule-related transcription factors, including Nodulation signaling pathway (NSP1 and NSP2), Ethylene response factor (ERF) required for nodulation (ERNs) and ERF required for nodule differentiation (EFD). These transcription factors enhance the expression of Nod factor-responsive genes by directly binding to the NF-box (Gleason *et al.*, 2006; Murray *et al.*, 2007; Tirichine *et al.*, 2007; Hirsch *et al.*, 2009). However, they are not distinct how DMI3 regulates nodule-related Nodule inception (NIN) and ERN, how EFD inhibits nodulation probably by disrupting cytokinin (Ryu *et al.*, 2012).

Nodulation is nitrogen fixation zone, Within the symbiosomes, the bacteria undergo a morphological terminal differentiation involving cell elongation and genome amplification, a process which is governed by numerous plant nodule-specific cysteine-rich peptides (NCR), which are addressed to the bacterium-containing compartments, where bacteroids reduce atmospheric nitrogen to ammonium. Oxygen-binding leghemoglobin functions to maintain a microaerobic environment necessary for bacteriod nitrogenase activity. Ammonia(um) is the main product of N_2_ fixation that is released from bacteroid and transported via ammonium transporter across the peribacteroid membrane (PBM) to the plant where initial assimilation into amino acids (AAs) occurs (White *et al.*, 2007 and 2009). Four key enzymes were activated for the primary assimilation of NH_4_^+^ in nodules. Glutamine synthetase (GS) and glutamate synthase (GOGAT) are collectively referred to as the GS-GOGAT pathway, conjunction with aspartate aminotransferase (AAT) and asparagine (ASN) synthetase (AS). Depending on the N-exporter nature, the determinate-nodule primarily transports allantoin (ALN) or/and allantoic acid (ALC) as fixed-N compounds (ureide-exporter), and the indeterminate-nodule export fixed-N predominantly as ASN and glutamine (GLN) (amide-exporter) (Sprent and James, 2007). Three *L. Japonicus* genes SST1 (symbiotic sulfate transporter), FEN1 (fail in enlargement of infected cells) and IGN1 (ineffective greenish nodules 1) and one *M. truncatula* gene, DNF1 (defective in nitrogen fixation), have been identified by analyses of Fix^-^ mutants. SST1 is a sulfate transporter localized in the peribacteroid membrance (PBM) and transfers SO_4_^2-^ from plant cytosol to bacteroids. FEN1 is homocitrate synthetase which supplies homocitrate to bacteroids to support synthesis of the nitrogenase complex. IGN1 is localized in the plasma membrane (PM) and plays important functions in symbiosome and/or bacteroid differentiation and maintenance. (Krusell *et al.*, 2005; Kumagai *et al.*, 2007; Hakoyama *et al.*, 2009; Kouchi *et al.*, 2010). The host plants control the differentiation of *rhizobia* into bacteria targeting nodule-specific cysteine-rich (NCR) peptides which direct symbiotic *rhizobia* into terminal bacteroid differentiation through the nodule-specific protein secretion system in indeterminate nodules (Wang *et al.*, 2010).

These studies in molecular genetics using two model legume plants, *Lotus japonicas* and *Medicago truncatula*, have identified a number of host genes thatessential for symbiotic nodule formation and nitrogen fixation activation. Soybean (*Glycine max* (L.) Merr.) is one of the most important legume crops for seed protein and oil content. In the past, soybean analysis of early gene expression in *rhizobia* symbioses focused on nodulin genes or only a few genes. These genetic loci (namely *Rj*/*rj*s) have been identified as controlling nodulation traits upon inoculation with compatible species (Hayashi *et al.*, 2012). *Rj* and/or *rj* genostypes were summarized into three types, as follows: (1) Recessive alleles at three loci, *rj1*, *rj5* and *rj6*, result in non-nodulation phenotypes (Williams and Lynch, 1954; Pracht *et al.*, 1993). (2) Recessive locus, Known as *rj7* or *nts1* (*nitrate-tolerant symbiosis*) causes a so-called ‘hypernodulation’ phenotype, the formation of an unusually large number of nodules (Caroll *et al.*, 1985; Akao and Kouchi, 1992). (3) Dominant alleles, *Rj2*, *Rj3*, *Rj4* and *Rfg1* have unique features that restrict nodulation with specific strains (or serogroups) of *Bradyrhizobium* or *Ensifer*/ *Sinorhizobium* (Cadwell, 1966; Vest and Caldwell, 1972; Weier *et al.*, 1990; Trase, 1995). These *Rj* and/or *rj* genes were map-based cloning using establishment of the resources for genomics studies of these model legumes and the soybean whole genomics sequencing (Schmutz *et al.*, 2010). For instance, *rj1* (*nod49*) and *rj5* (*nod139*), related lipo-oligochitin LysM-type receptor kinase genes, are characterized using positional cloning and candidate gene approaches genes (Indrasumunar *et al.*, 2010 and 2011). *Rj2* (*Rfg1*) are allelic genes encoding a member of the Toll-interleukin receptor/ nucleotide-binding site/leucine-rich repeat (TIR-NBS-LRR) class of plant resistance (R) proteins using map-based cloning (Yang *et al.*, 2010).

As functional genomics and sequence develop, studies of transcriptional profiles during nitrogen fixation are important to gain greater understanding of the related nitrogen fixation genes. Some studies using soybean evaluated nodulation gene-expression profiles in roots inoculated with *B.japonicum* and elucidated reduction of plant defenses (Brechenmacher *et al.*, 2008; Libault *et al.*, 2010; Carvalho *et al.*, 2013). The bacteroid differentiation and their nitrogen fixation are under strict control with complex interactions between the host legume cells and the intracellular bacteria; however, the mechanisms underlying differentiation of endosymbiotic *rhizobia* in symbiosomes to the bacteroid form are still largely unknown.

To improve our knowledge of genetic regulation during later differentiation nodulation and nitrogen fixation activation, we analyze gene expression of R1 stage (first flower) in isolated soybean root inoculated or mock-inoculated with *rhizobium*. This study aimed at analyzing the global expression change of genes in soybean roots. The results provide interesting insight into the evolution of genes specifically involved in the nitrogen fixation process.

## MATERIALS AND METHODS

### Bacterial strains and plant material

*Bradyrhizobium japonicum* USDA110 and *Sinorhizobium fredii* 45436 were grown at 28 □ and darkroom in liquid yeast extract mannitol broth (YMB) medium (pH 6.8) with moderate shaking (120 rpm). After 6 d, *B. japonicum* cells were amassed by centrifugation (4000 rpm, 10 min), washed with sterile water 3 times, and diluted in water to an optical density at OD_600_ = 0.8.

Soybean seeds of cultivar *Nannong 1138-2* were surface sterilized by soaking for 8 min in 0.1% HgCl_2_ and rinsed 5 times with sterile water. Seeds were germinated and grown in sterile ceramic pots containing sterile soil with 5 seedlings per pot under greenhouse conditions and natural light. The plants were inoculated at the V1 stage of development, by respectively inoculated inoculums containing USDA110, CCBAU45436 (50 ml of liquid suspension per pot at 0.8 OD_600_). Mock-inoculated plants (CK) received the same amount of autoclaved inoculums. Roots were harvested at R1 inoculated with USDA 110, CCBAU45436 and CK. Subsequently, the roots were extirpated nodulation and separated from shoots, immediately frozen in liquid nitrogen and stored at -80 □.

### RNA extraction, isolation of mRNA and cDNA synthesis

Total RNA was isolated from the roots of each treatment using Trizol reagent (Tiangen, Beijing), according to the manufacturer’s instructions. After extraction, the quality and quantity of the total RNA were analyzed by Nanodrop (Thermo Fisher Scientific Inc.) and Bioanalyser 2100 (Agilent Technologies). Pair-end index libraries were constructed according to the manufacture’s protocol (NEBNext^®^ Ultra^™^ RNA Library Prep Kit for Illumina^®^).

The isolation, fragmentation and priming of ploy(A)’s mRNA were performed using NEBNext Poly(A) mRNA Magnetic Issolation Module. The first and second strand cDNA were synthesized using ProtoScript □ Reverst Transcriptase and Second Strand Synthesis Enzyme Mix, respectively. The double-stranded cDNA was purified using AxyPrep Mag PCR Clean-up (Axygen) and treated with End Prep Enzyme Mix for end repairing. Then, the cDNA was added 5’ phosphorylation and dA-tailing in one reaction and was ligated to adaptors with a “T” base overhang.

### Sequencing and sequence aligment

Approximate 400 bp fragments of adaptor-ligated DNA (with the approximate insert size of 250 bp) were recovered using AxyPrep Mag PCR Clean-Up (Axygen). Two samples of USDA110 and CK were amplified by PCR for 11 cycles using P5 and P7 primers which can anneal with flow cell to perform bridge PCR and P7 primer carrying a six-base index allowing for multiplexing. The PCR productions were further cleaned up using AxyPrep Mag PCR Clean-up (Axygen), validated using an Agilent 2100 Bioanalyzer (Agilent Technologies), and quantified by Qubit and real time PCR (Applied Biosystems). Then libraries with different indexes were multiplexed and loaded on an Illumina HiSeq (Genewiz, Beijing) instrument according to manufacturer’s instructions (Illumina, San Diego, CA, USA). Sequencing was carried out using a 2 × 100 paired-end (PE) configuration; image analysis and base calling were conducted by the HiSeq Control Software (HCS) + OLB + GAPipeline-1.6 (Illumina) on the HiSeq instrument.

The reads from sequencing were aligned against the Williams 82 genome sequence (ftp://ftp.jgi-psf.org/pub/compgen/phytozome/v9.0/Athaliana/) using tophat-2.09 software, allowing a maximum of two mismatches.

### Bioinformatics analysis

A level of gene expression by expected number of fragments per kilobase of transcript sequence per million base pairs sequenced (FPKM value) was calculated using Rsem (v1.2.4) software. We calculated the differences in gene expression among different samples with EdgeR (v3.4.2) and *p*-value of significant difference. *P* value less than 0.05 was thought as significant difference.

GO enrichment analysis of differentially expressed genes using default GO association files was performed with “GO Term Finder” (http://go.princeton.edu/cgibin/GOTermFinder) where statistical significance (*p*-value) was calculated based on hypergeometric distribution with Bonferroni multiple testing correction and flase discovery rate (FDR) calculation as described. The GO enrichment analysis was performed with adjusted GO association files, Ontologizer, which (http://www.charite.de/ch/medgen/ontologizer/) was using “GO Term Finder” with a threshold *P* value of less than 0.01. The pathway annotation and enrichment of differentially expressed genes were performed using KAAS software with a threshold *P* value of FDR less than 0.05.

### Validation of RNA-Seq data by quantitative Real-time PCR (qRT-PCR) and results according with other *rhizobium*

Quantitative real-time PCR experiments were performed to validate the RNA-Seq results for 19 gene transcripts whose expression differed by more than 2.0-fold among USDA110, CCBAU45436 and CK. Primers for qRT-PCR were designed using Primer 3 software (http://frodo.wi.nit.edu/primers). Approximately 2 μg of purified total RNA were reverse transcribed using M-MLV reverse transcriptase (Invitrogen) with Oligo (dT)_20_ as primer (Invitrogen), according to the manufacturer’s instruction. Quantitative PCR was performed on an ABI ViiA^™^ 7 (Applied Biosystems) with the LightCycler system. PCR mixtures (final volume 20 μl) including 0.4 μl (approximately 50 ng) of first strand cDNAs, 0.5 μM each primer, 10μl of 2 × SYBR^®^ Green Realtime PCR Master Mix (TOYOBO). The cycling conditions were as follows: 2 min denaturation at 95 □ followed by 40 cycles of 95 °C for 10 s, 60 °C for 15 s, and 72 °C for 25 s. Thirteen candidate gene’s primers (Tab. 1) were designed according to every candidate gene sequence (http://soybase.org/gbrowse/cgi-bin/gbrowse/gmax1.01). Expression levels of these genes were normalized by tubulin (NCBI accession No. AY907703). Gene expression was quantified using the relative quantification (ΔΔC_T_) method and data was compared with internal controls. Each sample was replicated three times.

**Table 1.**
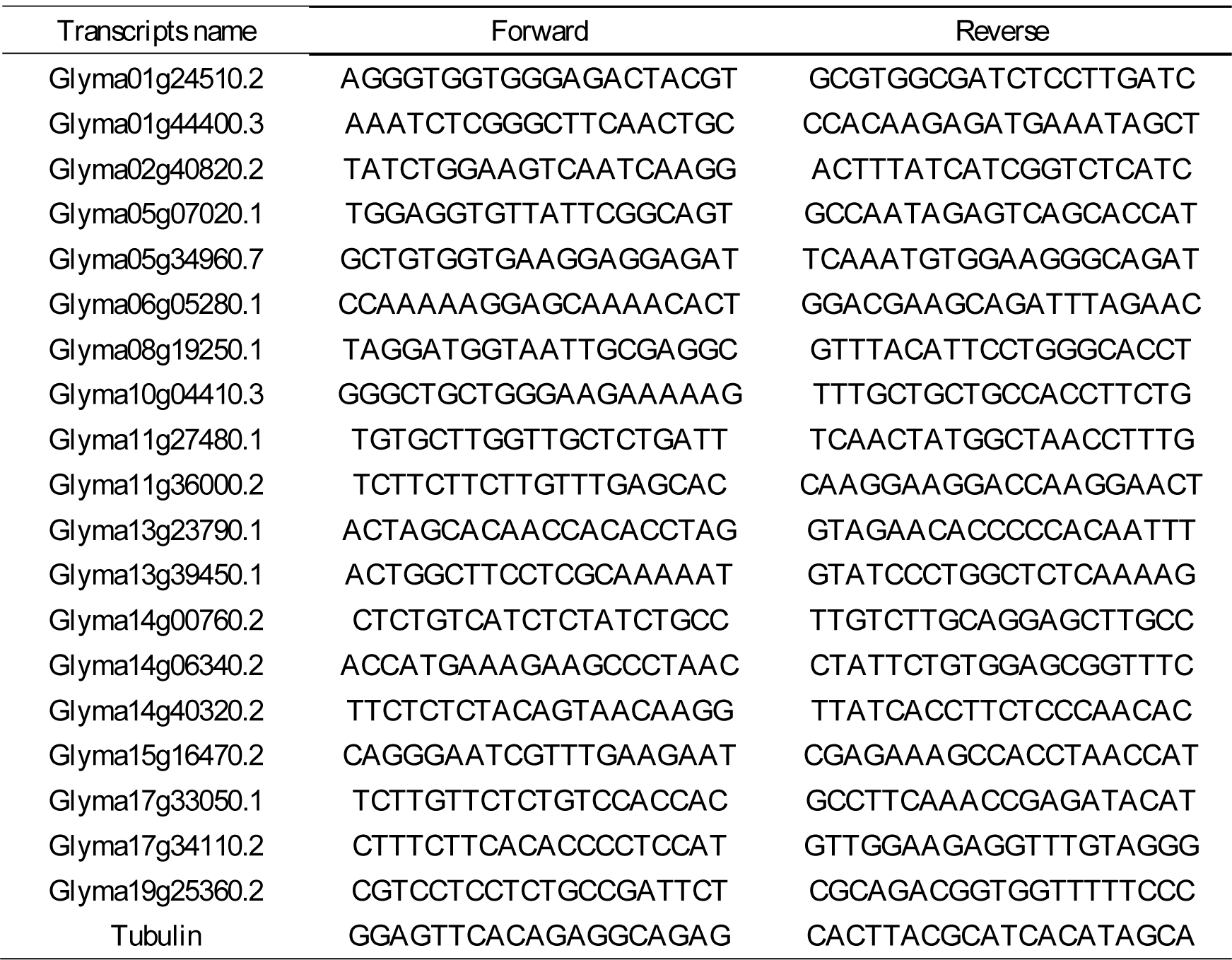
Primer sequences used for qRT-PCR

## RESULTS

### RNA-Sequencing analysis

Changes of plant roots’ transcript levels were analyzed between the inoculated USDA110 and CK at R1 stage by RNA-Seq. A total of 147.5 million reads were generated by 101 bp single-end sequencing from the two cDNA libraries, constituting 14.9 Gb of cDNA sequence (raw data). The raw data were filtrated using NGSQCToolkit (V2.3) according to the following processes: (1) The joint sequences were removed; (2) These reads were retained that these loci of the quality value higher than Q30 (Percentage of error probability less than 0.1%) accounted for 70% or more of total length of reads; (3) A base of 3’-end was removed; (4) These loci of the quality value lower Q30 were removed; (5) The reads of length lowing 70 bp were removed. Approximately 85.41% of the sequenced reads were high quality filtered reads (Tab. 2, Fig. S1). 92.46% of the clean data were successfully aligned to the soybean genome reference sequence (Glyma 1.01, http://www.phytozome/v9.0/athaliana) using Tophat-2.09 software. Thereinto, the reads of uniquely aligned to the reference genome (unique_mapped) accounted for 36.67% of total clean data (Tab. 3).

**Table 2.** Statistics of the Illumina HiSeq reads and comparison to the *G. max* reference genome (gmax1.01) Number is number of reads. Length is average length of reads. Q20 is percentage of error probability less than 1%. Q30 is percentage of error probability less than 0.1%. GC = (G + C) number/ total bases number × 100%. N is content of unidentifiable bases per million bases.

| Samples |  | Number | Length | Size of<br>bases (bp) | Q20(%) | Q30(%) | GC(%) | N(ppm) |
| --- | --- | --- | --- | --- | --- | --- | --- | --- |
| CK | Raw data | 71053298 | 101 | 7176383098 | 95 | 91 | 51 | 66 |
|  | Clean data | 60681336 | 100 | 6068133600 | 99 | 97 | 50 | 35 |
| USDA110 | Raw data | 76471968 | 101 | 7723668768 | 96 | 91 | 50 | 69 |
|  | Clean data | 65992290 | 100 | 6599229000 | 99 | 97 | 50 | 36 |

**Table 3.** Results of clean reads were aligned to reference genome Total_mapped is the total number of reads was aligned to reference genome. Unique_mapped is the number of reads which were aligned to exclusive position of reference genome. Muti_mapped is the number of reads which were aligned to multi-position of reference genome. PE_mapped is the number of reads of PE sequencing double end which were synchronously aligned to reference genome.

| Samples | Total_mapped | Unique_mapped | Multi_mapped | PE_mapped | Mapped percent(%) |
| --- | --- | --- | --- | --- | --- |
| CK | 55793166 | 20436743 | 35356423 | 27327800 | 91.94 |
| USDA110 | 61369446 | 22522473 | 38846973 | 30114865 | 92.99 |

### The differentially expressed genes profiles responding to *rhizobium* inoculation

A total of 55787 transcripts were quantified based on FPKM value of unique_mapped fragments for the analysis of gene expression (Tab. S1). There are 596 differentially expressed transcripts (P value ≤0.05 and fold-change ≥ 2.0) by edgeR program of Bioconductor software through FPKM values (Tab. S2). Of these, 280 transcripts were up-regulated and 316 were down-regulated in inoculated USDA110 VS CK (Fig. 1). The most log_2_^CPM^ value of transcripts expression level were less than 5.0, the correlation relation showed as figure 2 between transcripts expression level and expression difference.

**Figure 1.**
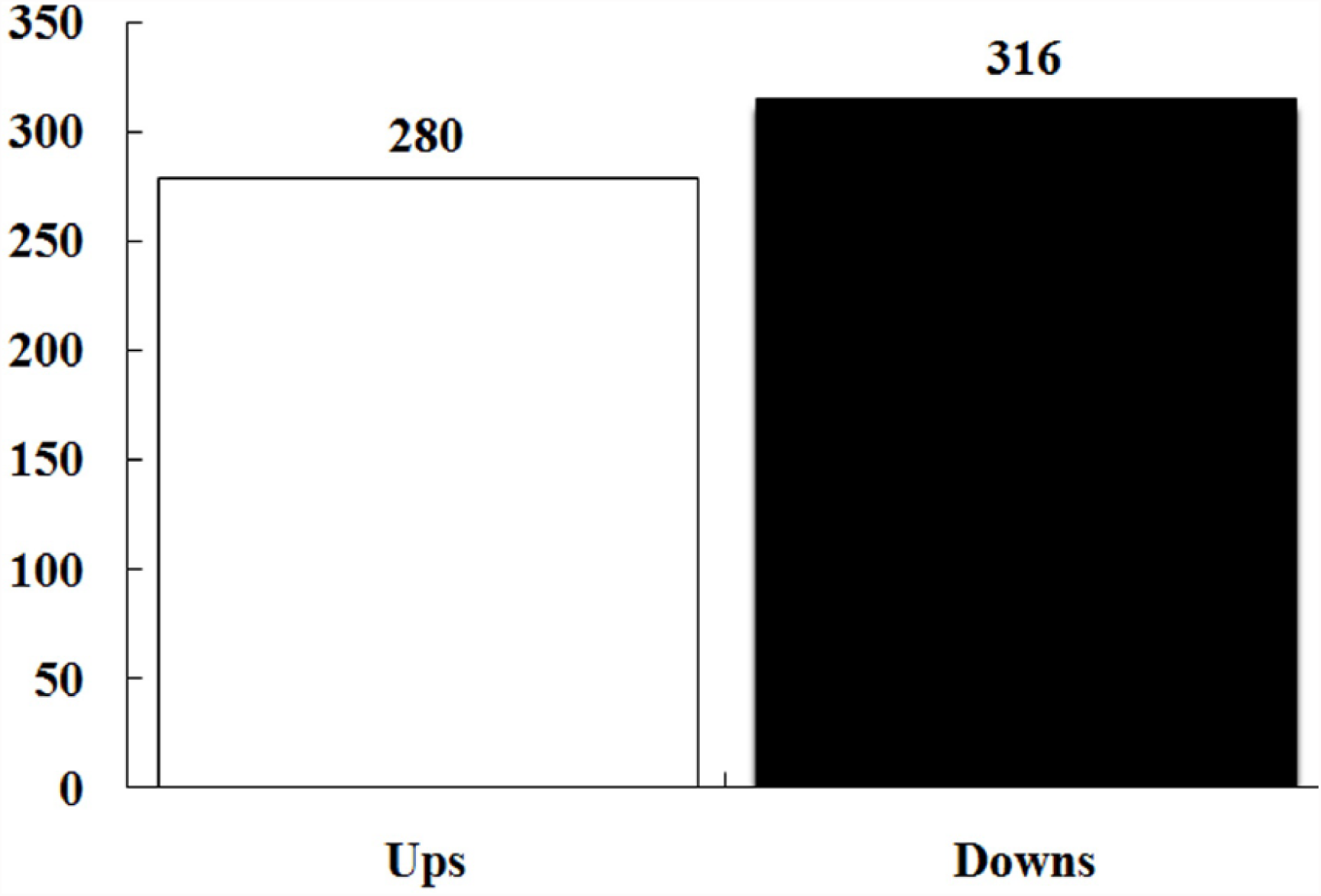
Number of differentially expressed transcripts in CK VS USDA110.

**Figure 2.**
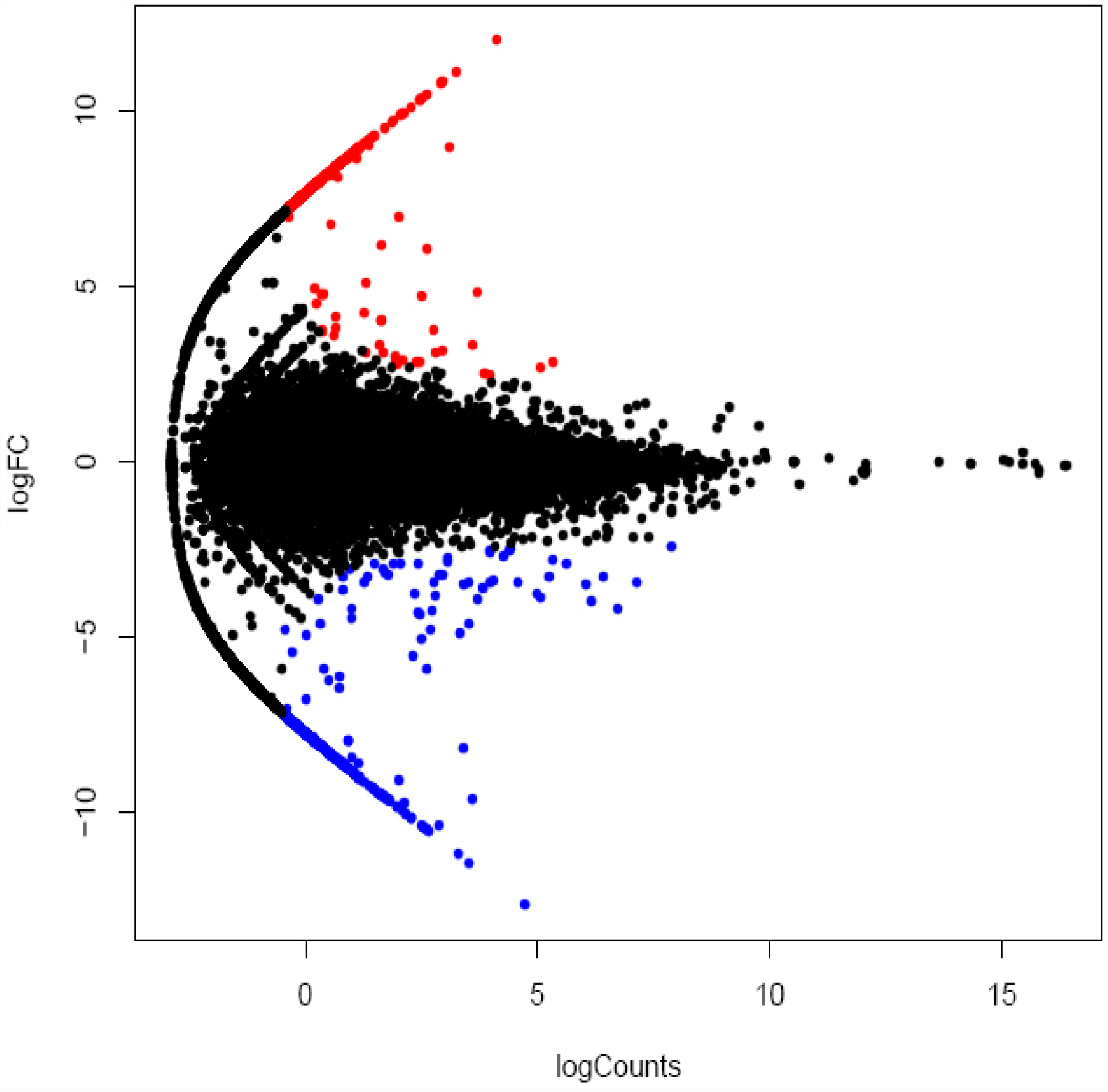
Relation between transcripts expression difference and counts log^FC^ is value of log2^fold change of transcript expression^. log^CPM^ is average value log2^counts per million reads of transcript expression^. Red dot represent up-regulated transcripts. Blue dot represent down-regulated transcripts. Black dot represent no significantly different transcripts.

### Gene ontology category analysis

As a useful tool for gene functional annotation, WEGO (Web Gene Ontology Annotation Plot) has been widely used in many soybean studies (Li *et al.*, 2013). It has become one of the most tools for downstream gene annotation analysis studies. In this research, GO assignments were used to classify the functions of the predicted nitrogen fixation genes. A total of 39163 expressed genes of two materials were converted into GO-identities (IDs) (P value ≤0.01) and classified three functional categories; cellular component, molecular function and biological process by mapping to GO Term Finder (http://search.cpan.org/dist/GO-TermFinder/lib/GO/TermFinder.pm) (Fig. 3). The differentially expressed genes (DEGs) (464) according to three categories with a gene ontology annotation were further classified into subsets. There were 5 subsets within the cellular component category, 3 subsets within the molecular function category, and 11 subsets within the biological process category. The genes were classified as follows: 104 genes mapped to cell component category; 182 genes mapped to molecular function category; and 178 genes mapped to biological process category. Five subsets of cellular component category are cell or cell part, organelle or organelle part, membrane-enclosed, membrane or membrane part, and non-membrane-bounded organelle. Three subsets of molecular function category are catalytic activity, binding, transporter, and binding. Eleven subsets of biological process category are metabolic process, biological regulation, cellular component organization or biogenesis, celluar process, reproduction, multicellular organismal process, signaling, developmental process, single-organism process, response to stimulus, and localization (Fig. 4 and supplementary dataset S1). The most abundant unigenes were related to cell or cell parts in cellular component category, catalytic activity and binding in molecular function category, and metabolic process, response to stimulus and localization in biological process category. The results showed that symbiotic fixation nitrogen brought on a series of plant’s genes changes.

**Figure 3.**
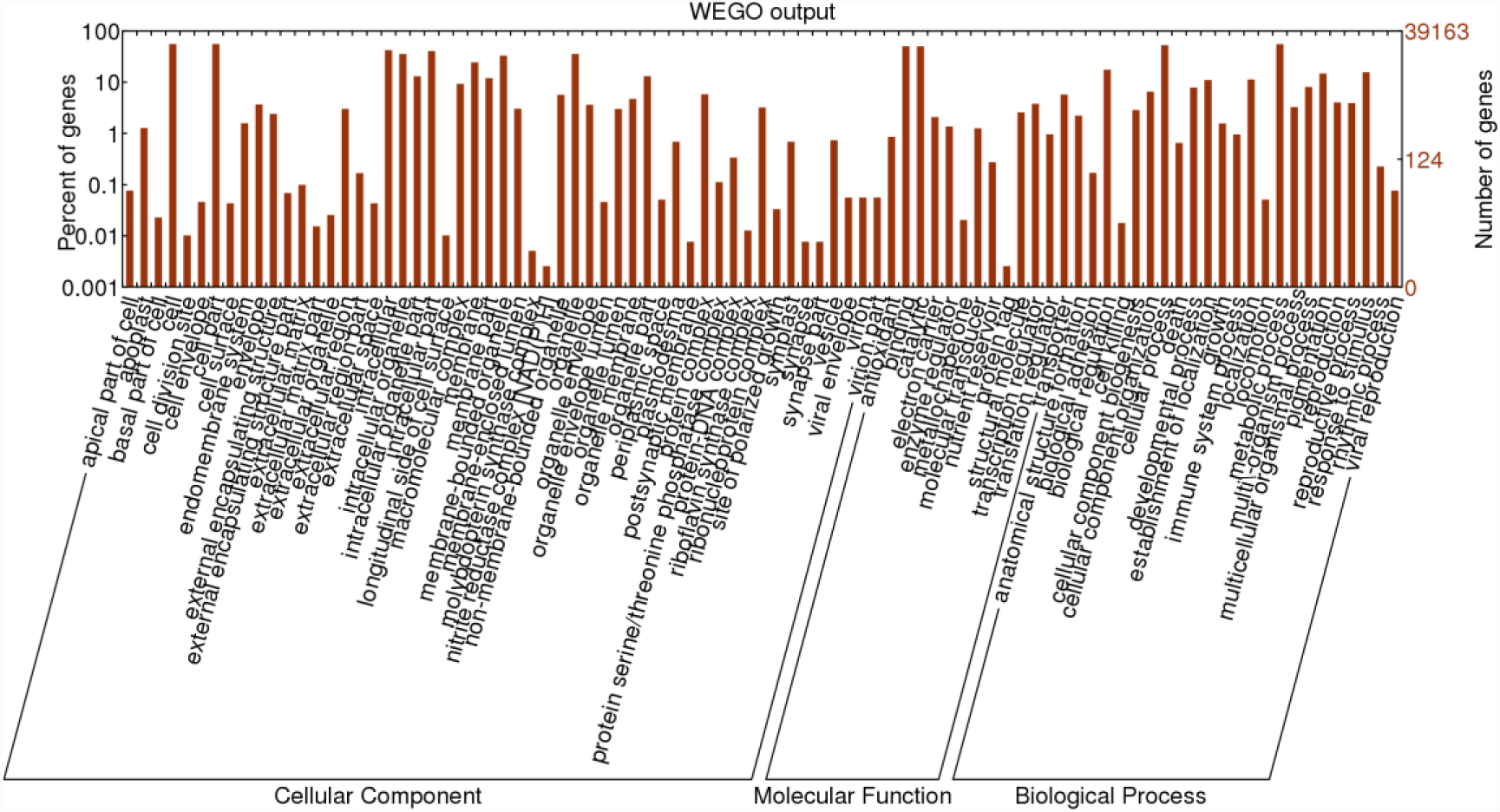
Functional categorization of expressed genes in plant roots of inoculated CK and USDA110.

**Figure 4.**
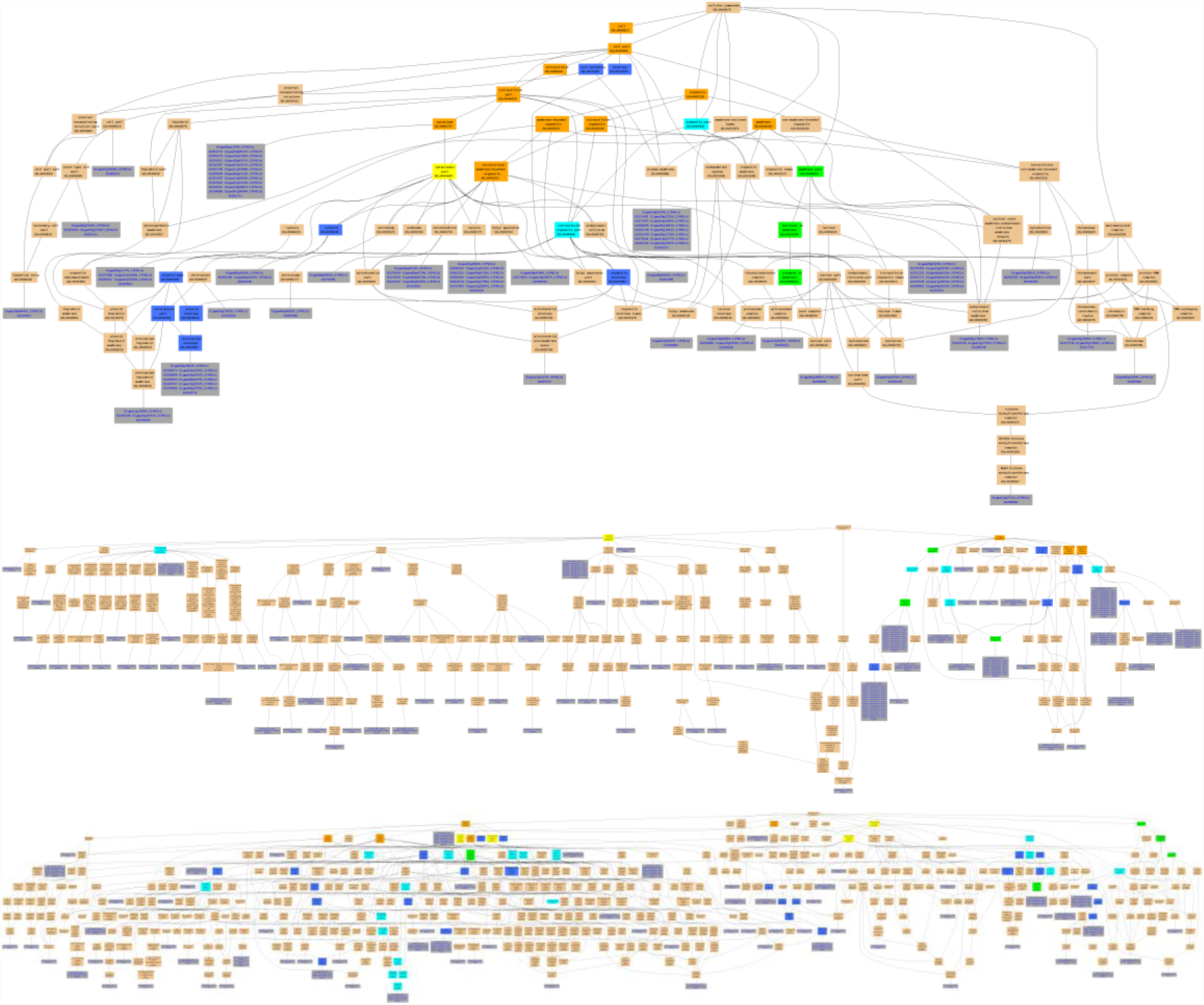
Cellular Component, Molecular Function and Biological Process of differentially expressed genes between CK and USDA110.

### Pathway enrichment analysis of DEGs

Genes usually interact with each other to carry out certain biological functions. Knowledge of the expressions of multiple genes and their regulation in symbiotic fixation nitrogen biosynthesis is required to further understand the regulatory mechanisms. Pathway-based analysis helps to clarify the biological functions of genes and signal transduction pathways associated with DEGs compared with the whole genome background. For our research, 169 biological pathways were identified by Kyoto Encyclopedia of Genes and Genomes (KEGG) pathway analysis. There are 40 pathways related to carbon cycle and metabolism, 16 pathways related to amino acid synthesis and metabolism, 8 pathways related to hormone, 8 pathways related to signaling pathway, 6 pathways related to vitamin, 3 pathways related to alkaloid, and related to RNA and virus etc. Thereinto, a nitrogen metabolism and a NOD-like receptor (NLR) signaling pathway were identified. These websites of pathway’s map were also listed in table S3. A total of 98 difference expressed genes (115 transcripts) with pathway annotations were identified by KAAS software. The expressed level of 54 transcripts was up-regulated, and 61 transcripts were down-regulated. The number of transcript associated with pathway ranges from 1 to 44, with an average number of 4.38 pathways per transcript. 23 transcripts were associated with one pathway, 92 transcripts were associated with two or more pathway. The transcripts of Glyma10g04410.2 was involved in 44 pathways, secondly, Glyma14g00760.2 was involved in 17 pathways (Tab. S4).

From those pathways, we selected the NOD-like receptor signaling pathway (Fig. 5) and nitrogen metabolism pathway (Fig. 6) for further analysis. There is only one up-regulated transcript (Glyma14g40320.4) in the NLR signaling pathway. Seven differentially expressed transcripts associated with the nitrogen metabolism pathway. The all 7 transcripts were down-regulated, which may be because symbiosis provided nitrogen nutrition for plant, some nitrogen transformation genes needn’t over-expressed.

**Figure 5.**
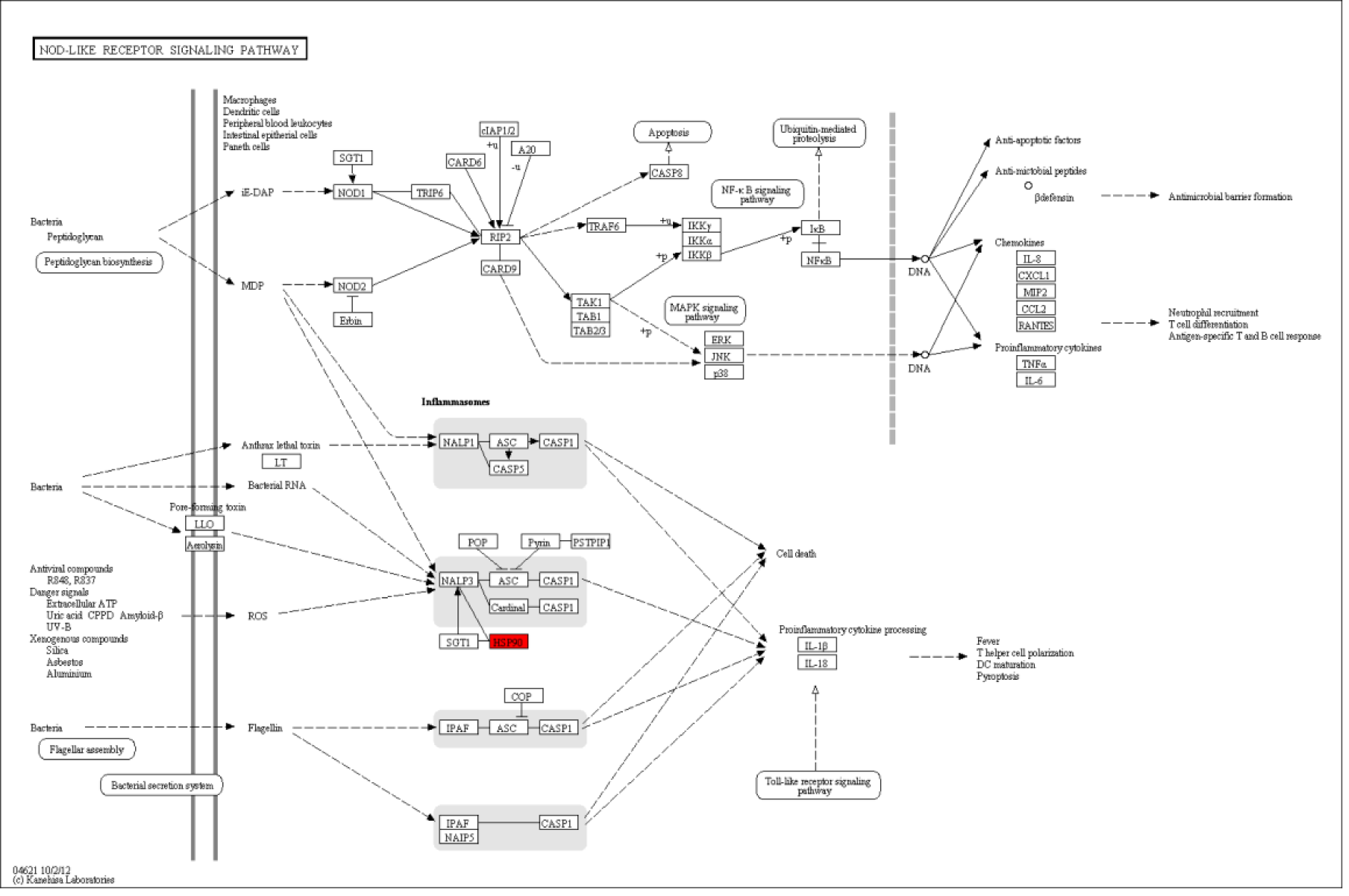
NOD-like receptor signaling pathway. Note: Red rectangles indicate up-regulated genes. Green rectangles indicate down-regulated genes. The same below

**Figure 6.**
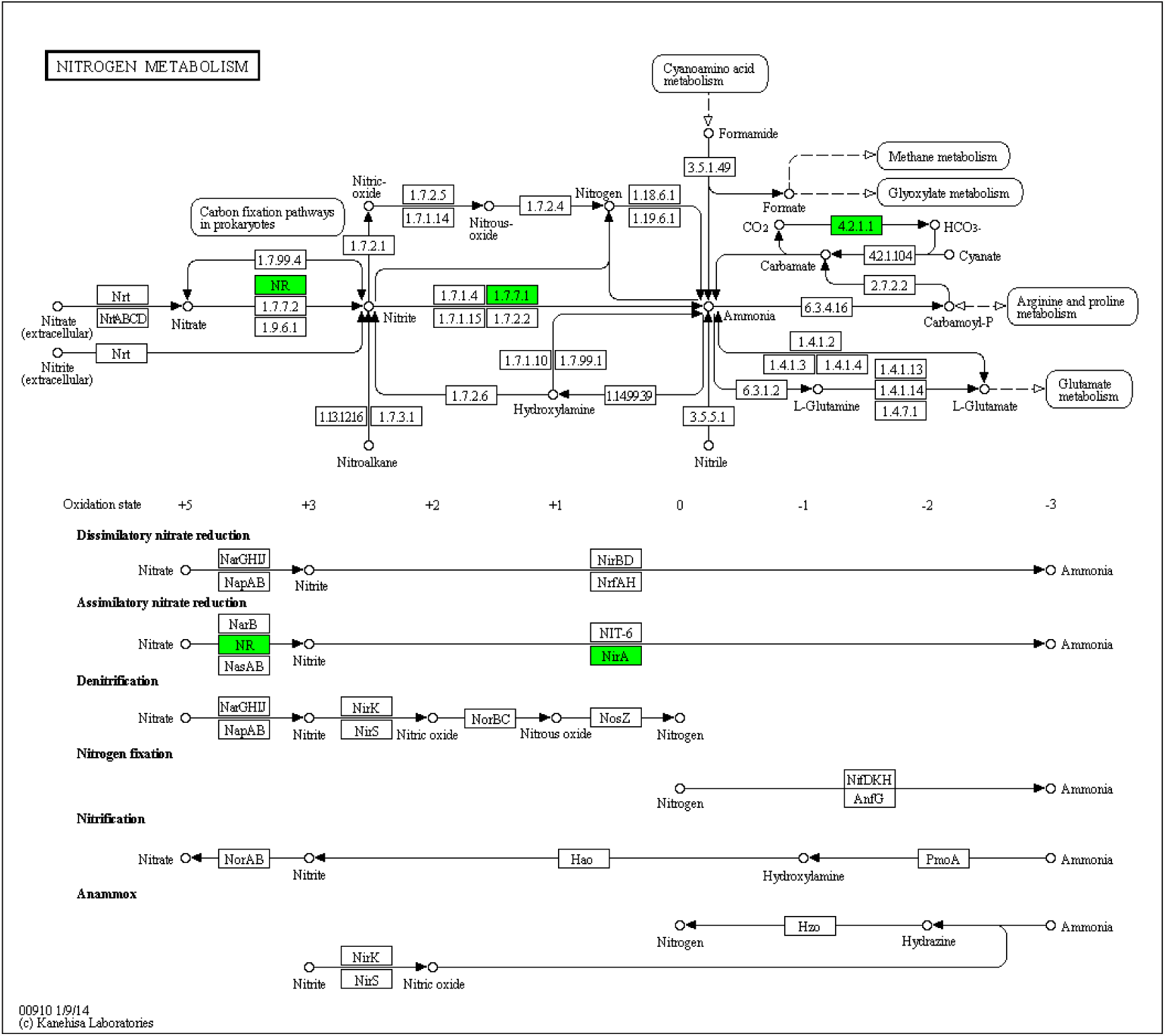
Nitrogen metabolism pathway.

### Validation of RNA-Seq data by qRT-RCR and result’s university in other *rhizobium*

To confirm the expression patterns validity by RNA-Seq in the soybeans of mock-inoculated, inoculated *B. rhizobium* USDA110 and result’s university in other *rhizobium*, we used qRT-PCR analyses to analyze expressions of 19 candidate genes of CK, USDA110 and CCBAU45436. Results showed significant differences in the expression levels between the inoculated and mock-inoculated *rhizobium* for 19 transcripts (Fig. 7). Ten transcripts (Glyma01g44400.3, Glyma02g40820.2, Glyma08g19250.1, Glyma11g36000.2, Glyma13g39450.1, Glyma14g40320.2, Glyma15g16470.2, Glyma17g33050.1, Glyma17g34110.2 and Glyma19g25360.2) were significantly up-regulated in soybean roots of inoculated *rhizobium*, other nine transcripts were significantly down-regulated. Although the RNA-Seq values showed slight variations compared with the corresponding values from the qRT-PCR analyses, the tendency of expression level from RNA-Seq analysis was consistent with those obtained by qRT-PCR. The expression levels of 19 transcripts in soybean roots inoculated USDA110 strain were similar to inoculated CCBAU45436 strain. These results highlighted the fidelity and reproducibility of the RNA-Seq analysis used in the present study. The results had a certain extent’s university in different *rhizobium* strains.

**Figure 7.**
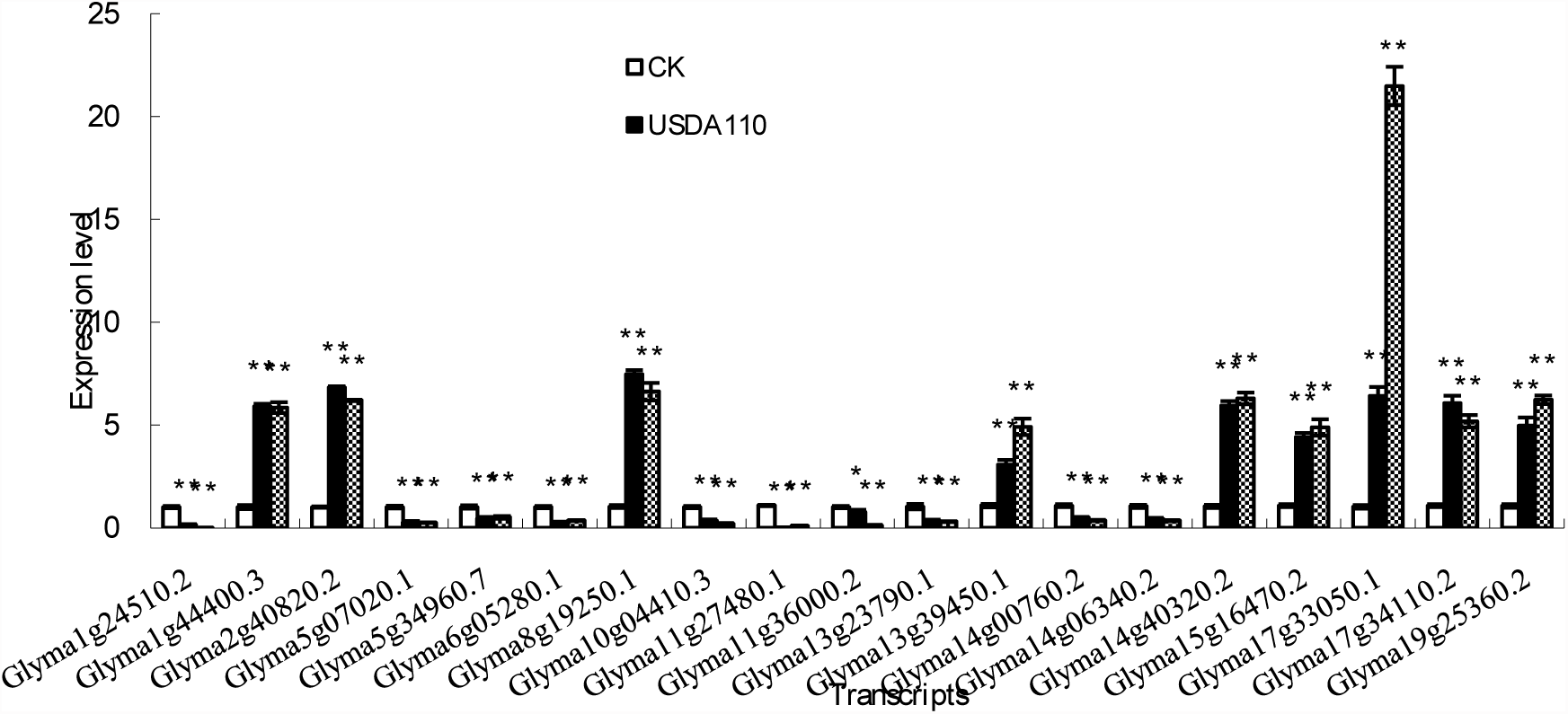
Analysis of 19 differentially expressed genes using RNA-Seq by qRT-PCR in CK, USDA110 and CCBAU45436.

## DISCUSSION

The role of biological nitrogen fixation (BNF) in world crop production has been emphasized for several decades. The molecular mechanisms involved in the interaction between legumes and *rhizobia* has greatly improved (Kouchi *et al.*, 2010), specifically expressed during the symbiotic nodule formation (Libault *et al.*, 2010; Hayashi *et al.*, 2012). The development of nitrogen-fixing nodules on roots of leguminous plants commences with a molecular dialogue between the host plant and a compatible strain of *rhizobia*. The identification of plants that spontaneously form nodules, together with the observations that ectopic application of cytokinins or auxin transport inhibitors to the root surface lead to the development of nodule-like structures, demonstrates that the machinery required for nodule primordium (NP) development is intrinsic to the plant (Gleason *et al.*, 2006; Murray *et al.*, 2007; Tirichine *et al.*, 2007; Van *et al.*, 2015). More recently, comprehensive analysis by means of proteomics and transcriptome have revealed that more than a thousand genes are specifically induced or highly enhanced in nodules (Wienkoop and Saalbach, 2003; Høgslund *et al.*, 2009; Carvalho *et al.*, 2013). However, few progresses have been achieved towards host plant to control bacteroid differentiation and activating *rhizobial* nitrogen fixation.

This study has provided a new data set identifying the expression of DEGs during symbiotic nitrogen fixation in R1 stage. Massive parallel sequencing identified 596 differentially expressed transcripts from two contrasting libraries. We examined gene expression levels in detail, and found significant difference between inoculated and mock-inoculated *rhizobia*. Among the differentially expressed genes, more were down-regulated than up-regulated, and some showed a differential expression pattern in all three contrast groups, indicating that there were overlaps at the transcriptional level. The fact that there were many down-regulated transcripts indicates that there are more negatively regulated genes than positively regulated ones with functions in the nitrogen fixation pathway. To explore the genes with unknown functions, the expression patterns of different genes were analyzed by according to similarities in expression profiles.

GO analysis and the pathway enrichment analysis of DEGs showed that most are related to carbon and amino acid synthesis or metabolism. In the symbiotic fixation nitrogen, NOD-like receptor signaling pathway, the nitrogen metabolism and plant hormone play an important role. The Hsp90 of host plant was up-regulated by inoculated *rhizobium*. Hsp90 is an abundant, dimeric ATP-dependent molecular chaperone, and ATPase activity is essential for its functions (Retzlaff *et al.*, 2009). The NLRs are classified as part of the signal transduction ATPases with numerous domains (STAND) clade within the AAA+ ATPase family (Proell *et al.*, 2013). The interaction of heat shock protein 90 (Hsp90) and suppressor of the G2 allele of Skp1 (SGT1) activate Nod1 (Correia *et al.*, 2007). The nodulation ability of host plants could be improved by boosting up expression’s level of Hsp90.

In nitrogen metabolism, the nitrate reductase (NAD(P)H), ferredoxin-nitrite reductase and carbonic anhydrase of host plant were down-regulated by inoculated *rhizobium*. The enzyme nitrate reductase, ferredoxin-nitrite reductase and carbonic anhydrases respectively catalyzes the reduction of nitrate to nitrite (Okamoto *et al.*, 1993), the six-electron reduction of nitrite (oxidation state +3) to ammonium (oxidation state -3) in the second step of the nitrate assimilation pathway (Takahashi *et al.*, 2001) and the reversible hydration of CO_2_ to form HCO_3_^-^ and protons (Breton, 2001). The mostly ammonium demand of host plant could be basically satisified by the nitrogen fixation ability of *rhizobium*, fewer nitrate nitrogen was absorbed from soil, so three reductases’ activity decreased.

The plant hormone regulates a number of developmental and physiological processes including nodulation. For example, a cytokinin receptor triggers spontaneous root nodule organogenesis (Murray *et al.*, 2007; Tirichine *et al.*, 2007); *Rhizobia* and external stress signals activate mitogen activated protein kinase (MAPK) signaling cascades and the action of plant hormones including ethylene, salicylic acid (SA), abscisic acid (ABA), and jasmine acid (JA) which were negatively regulated nodulation (Ryu *et al.*, 2012). In this study, jasmonate ZIM domain-containing protein (JAZ) was down-regulated by inoculated *rhizobium* (Tab. S3). The result is consistent with previous study.

The validation results of qPCR were consistent with the transcripts expression patterns identified by RNA-Seq and expression levels of transcripts in soybean roots inoculated USDA110 strain were similar to inoculated CCBAU45436 strain. These showed that the results of RNA-Seq have a certain extent of universality in different *rhizobium* strains.

Previously, QTL mapping was used to locate biological nitrogen fixation traits in soybean along with 16 gene/QTL regions of 12 chromosomes (Madsen *et al.*, 2003; Searle *et al.*, 2003; Tanya *et al.*, 2005; Nicolás *et al.*, 2006; Yang *et al.*, 2010; Hayashi *et al.*, 2012; Santos *et al.*, 2013) (Tab. S5). There are significantly different genes of using RNA-Seq in these QTL regions. For example, three significant different transcripts (Glyma11g08720.2, Glyma11g09250.2 and Glyma11g09630.2) between inoculated and mock-inoculated *rhizobium* were located between Satt509 and Satt251 on chromosome 11 in study of *Santos* (2013). However, the *Rj*/*rj* genes involved in nitrogen-fixation root nodule formation in soybean weren’t detected in the study. These results may be because that the difference of nodulation gene wasn’t significant difference between inoculated and mock-inoculated *rhizobia* in soybean’s R1 stage. More novel genes were identified by RNA-Seq in the study and weren’t located by previous gene/QTLs mapping study. Their function needs to be further verified using other methods.

In the study, 280 and 316 transcripts were up-and down-regulated in R1 stage of soybean, respectively. Gene Ontology analyses detected 5, 3 and 11 subsets within the cellular component, molecular function and biological process category, respectively. A total of 169 biological pathways were identified by KEGG pathway analysis. Putative functions for some of these genes were assigned for the first time in the *rhizobium*-soybean symbiosis. Novel genes were firstly described and could be related to the nitrogen fixation process.

## ACKNOWLEDGMENTS

The authors are gratefully indebted to Prof. C. F. Tian, China Agricultural University, for donated *rhizobia* strains. This work was supported by Anhui Provincial College Program for Natural Science (KJ2013A077), Science and Technology Support Program of Sichuan Province (2015NZ0046) and Key Disciplines of Anhui Science and Technology University (AKZDXK2015B02).

## Legends to Supplementary Tables, Figures and Supplement dataset S1

Table S1 Expressed level of transcripts aligned to soybean genomic sequence GeneLength is length of reference geness. Sample_reads is number of gene’s exclusive alignment fragment. Sample_FPKM is gene’s expected number of fragments per kilobase of transcript sequence per million base pairs sequenced. The same below

Table S2 Differentially expressed transcripts of roots between CK and USDA110 log^FC^ is value of log_2_^fold change of transcript expression^. log^CPM^ is average value log_2_^counts per million reads of transcript expression^. PValue is P value of significantly different gene. FDR is false discovery rate.

Table S3 The KEGG pathway of differentially expressed genes CK-VS-USDA110 is differentially expressed gene’s number of two samples in the pathway. All-Diffgene is all genes’ number of gene’s background in the pathway. Genes are differentially expressed genes in the pathway. KOs are terms of KEGG Orthology. PathwayURL is websites of pathway’s map.

Table S4 Number of pathway of differentially expressed genes

Table S5 Previously genes/QTLs regions exist in differentially expressed genes

Note: NDW is nodule dry weight. SDW is shoot dry weight. NN is nodule number. NFW is nodule fresh weight, PDW is plant dry weight. NDW/NN is ratio nodule dry weight/NN.

Figure S1 Summary of quality check and filtering

Supplement dataset S1 GO analysis of differentially expressed genes

